# Transcriptomics in serum and culture medium reveal shared and differential gene regulation in pathogenic and commensal *Streptococcus suis*

**DOI:** 10.1101/2022.10.16.512421

**Authors:** Simen Fredriksen, Suzanne D. E. Ruijten, Gemma G. R. Murray, Maria Juanpere-Borràs, Peter van Baarlen, Jos Boekhorst, Jerry M. Wells

**Affiliations:** Host-Microbe Interactomics Group, Animal Sciences Department, Wageningen University, Wageningen, the Netherlands; Department of Veterinary Medicine, University of Cambridge, Cambridge, United Kingdom

## Abstract

*Streptococcus suis* colonizes the upper respiratory tract of healthy pigs at high abundance but can also cause opportunistic respiratory and systemic disease. Disease-associated *S. suis* reference strains are well studied, but less is known about commensal lineages. It is not known what mechanisms enable some *S. suis* lineages to cause disease while others persist as commensal colonizers, or to what extent gene expression in disease-associated and commensal lineages diverge. In this study we compared the transcriptomes of 21 *S. suis* strains grown in active porcine serum and Todd-Hewitt yeast broth. These strains included both commensal and pathogenic strains, including several strains of sequence type (ST) 1, which is responsible for most cases of human disease and considered the most pathogenic *S. suis* lineage. We sampled the strains during their exponential growth phase and mapped RNA-sequencing reads to the corresponding strain genomes. We found that the transcriptomes of pathogenic and commensal strains with large genomic divergence were unexpectedly conserved when grown in active porcine serum, but that regulation and expression of key pathways varied. Notably, we observed strong variation of expression across media of genes involved in capsule production in pathogens, and of the agmatine deiminase system in commensals. ST1 strains displayed large differences in gene expression between the two media compared to strains from other clades. Their capacity to regulate gene expression across different environmental conditions may be key to their success as zoonotic pathogens.

## Introduction

*Streptococcus suis* is an opportunistic pathogen that can cause septicaemia and meningitis in pigs and humans, but also colonizes the upper respiratory tract of healthy pigs [1]. Different *S. suis* lineages appear to be specialized to different niches, being made up of either predominantly clinical strains isolated from necropsy (pathogenic clades), or non-clinical strains isolated from the oral cavity of pigs (commensal clades) [2]. Clinical and non-clinical strains are, however, found in all clades, showing that pathogenic lineages can colonize the oral cavity of asymptomatic pigs at low abundance and that commensal lineages can invade the host in certain circumstances. Recent microbiome studies have confirmed that commensal *S. suis* colonize the oral cavity of piglets at high abundance [3–5].

Limited research has been conducted on commensal *S. suis* and the mechanisms underlying their great success as piglet oral biofilm colonizers. Commensal *S. suis* clades have high genomic diversity and larger genome sizes compared to pathogenic clades [6], and production of secondary metabolites may aid them in antagonising con- and heterospecific competitors [7]. Pathogens have conserved virulence-associated genes but reduced accessory genomes [6]. To successfully infect hosts, pathogens need to rapidly respond to different environments and shifts in nutrient availability and host immune responses. Firstly, they need to colonize the oral cavity to gain access to the host tissue and to persist in herds between outbreaks to ensure transmission. Secondly, they need to cross host epithelium, enter the bloodstream, and survive and proliferate in host tissues and body fluids with varying nutrient availability while avoiding eradication by host defences. To cause meningitis, *S. suis* also crosses the blood-brain barrier. *S. suis* niche differentiation is poorly understood, and the exact combination of virulence-associated genes and gene-host-environment interactions necessary to cause invasive disease have not been established. It is possible that regulation of gene transcription by phase variation and master switches such as carbon catabolite repression are key to rapidly adapting to a changing environment [8–10].

Despite the great heterogeneity of *S. suis* lineages, most research has been focused on a few closely related and highly pathogenic strains. Transcriptomic studies have used sequence type 1 (serotype 2) strains associated with zoonotic disease, such as P1/7 and S10 [11–16]. This has led to knowledge of how the most common pathogenic S. *suis* clade adapts to the host but left other pathogenic and commensal clades understudied. It is not known how well results from the commonly studied strains translate to other clades especially given the high functional redundancy of many virulence factors described for S. *suis* [17, 18]. A greater understanding of the species-wide S. *suis* transcriptome may yield insight into the differences between commensal and pathogenic S. *suis* clades and increase our understanding of S. *suis* ecology and evolution.

In this study we compare the transcriptomes of 21 S. *suis* strains from a wide phylogenetic background and different isolation sources, including not only clinical strains from pathogenic clades and non-clinical strains from commensal clades, but also clinical strains from commensal clades and non-clinical strains from pathogenic clades. We determined growth curves in Todd-Hewitt Yeast Broth (THY) and active porcine serum (APS), which contains complement and scarce amounts of essential metals [12], and extracted RNA during the exponential growth phase. We mapped RNA-seq data to the individual strain genomes and compared normalized sequencing coverage per gene. The resulting dataset allowed us to gain a better understanding of shared and niche-specific gene expression in commensal and pathogenic S. suis.

## Methods

### Strain selection

We selected 21 S. *suis* strains that are broadly representative of the species, including both clinical and non-clinical strains from different phylogenetic backgrounds (Table 1). Where available we included closely related clinical and non-clinical strains for comparison (i.e., not only clinical strains from pathogenic clades and non-clinical strains from commensal clades, but also clinical strains from commensal clades and non-clinical strains from pathogenic clades). While the isolation source and phylogenetic clade of a strain gives an indication of its virulent potential, this has only been experimentally tested *in vivo* for few strains. Strains P1/7 and S10 are known to be from a pathogenic clade and highly virulent, while strain T15 has been experimentally shown to have low virulence in pigs despite being from a pathogenic clade [19].

**Table 1:** Strain overview. Shortened 3-character name used in figures and tables, full original strain name, serotype, sequence type (ST), number of plasmids in genome assembly, and strain metadata. NT = non-typeable.

| Name | Strain | Country | Serotype | ST | Clade type | Isolation source | Plasmids |
| --- | --- | --- | --- | --- | --- | --- | --- |
| C01 | M101999_C1 | Spain | 8 | NT | Pathogenic | Non-Clinical | 1 |
| C15 | SS15055_N2_C15 | Spain | 2 | 1 | Pathogenic | Non-Clinical | 0 |
| D11 | DNS11 | Denmark | 9 | 16 | Pathogenic | Systemic/Brain | 1 |
| D13 | DNC13 | Denmark | NT | NT | Commensal | Non-Clinical | 2 |
| D15 | DNC15 | Denmark | 16 | NT | Commensal | Non-Clinical | 0 |
| D20 | DNS20 | Denmark | NT | NT | Commensal | Systemic/Brain | 2 |
| D43 | DNR43 | Denmark | 2 | 28 | Pathogenic | Respiratory | 1 |
| D48 | DNR48 | Denmark | 8 | NT | Pathogenic | Respiratory | 1 |
| D49 | DNC49 | Denmark | 2 | 28 | Pathogenic | Non-Clinical | 0 |
| G69 | DE609B | Germany | 2 | 1 | Pathogenic | Systemic/Brain | 0 |
| I12 | 21435_1 | Spain | 31 | NT | Commensal | Non-Clinical | 0 |
| I27 | 21437_3 | Spain | 4 | NT | Commensal | Non-Clinical | 0 |
| L42 | LSS42 | UK | 16 | NT | Pathogenic | Non-Clinical | 0 |
| P17 | P1/7 | UK | 2 | 1 | Pathogenic | Systemic/Brain | 0 |
| S10 | S10 | Netherlands | 2 | 1 | Pathogenic | Systemic/Brain | 0 |
| S11 | M102942_S11 | Spain | 9 | 123 | Pathogenic | Systemic/Brain | 1 |
| S20 | M104300_S20 | Spain | 2 | 1 | Pathogenic | Systemic/Brain | 0 |
| S26 | M105052_S26 | Spain | 19 | NT | Commensal | Systemic/Brain | 0 |
| S40 | M106471_S40 1 | Spain | 30 | NT | Commensal | Systemic/Brain | 0 |
| T15 | T15 | Netherlands | 2 | 19 | Pathogenic | Non-Clinical | 2 |
| X15 | 1521251 | Canada | 7 | NT | Pathogenic | Systemic/Brain | 0 |

This study aimed to assess the overall species transcriptome and compare groups of strains rather than focus on specific gene variants or individual strains. Thus, we prioritized RNA-sequencing of a single sample from many different strains (biological replicates) rather than technical replicates of each strain. To benchmark the repeatability of our methods we performed RNA sequencing on 3 technical replicates of S10 grown in each medium, processed separately on different days.

### Hybrid genome assembly

We created new Illumina-Nanopore hybrid genome assemblies for 19 of the 21 strains, as only strains P1/7 and S10 had complete genomes available. T15 had a circular genome assembly that lacked plasmids. We isolated DNA from overnight cultures using the PowerSoil DNA Isolation Kit (Qiagen). Generation of short-read sequencing data for the strains from Germany, Canada, and the UK has been previously reported [2, 3, 20]. Strains without available short-read sequencing data were 250 bp paired-end sequenced using an Illumina HiSeq 2500 instrument by MicrobesNG (Birmingham, UK). Nanopore sequencing was done with the SQK-LSK-109 Ligation Sequencing Kit and Guppy Software v5.0.16 with high accuracy base calling. Genomes were assembled using Unicyler v0.4.9 [21] with default settings. See table S1 for genome statistics.

### RNA extraction and sequencing

We sampled the strains in the mid exponential growth phase during growth in Todd-Hewitt Yeast Broth (THY) and active porcine serum (APS). THY is a rich lab medium used to rapidly grow streptococci to high density, and consists of several ingredients including meat infusion, tryptone, glucose, and yeast extract. Serum is extracted from blood by removing cells and clotting factors and although it contains high glucose concentrations essential minerals are scarce. For instance, iron is sequestered by host transferrin to limit bacterial growth in blood. Active serum is not heat-treated and may thus contain complement factors and antibodies binding to S. *suis* as this pathogen is endemic on farms. S. *suis* has several mechanisms to evade complement activation and formation of the membrane attack complex, including the capsule, factor H binding proteins, and other less well understood mechanisms [17, 22].

Growth curves were started with an overnight culture of *S. suis* in THY and adjusted to OD_600_ 0.5 by dilution in PBS and subsequently inoculated 1:10 in either THY or APS. For determination of growth curves the strains were grown in 96-well plates with 200 μL total volume per well and incubated at 37 °C with 5% CO_2_. The plates were shaken to prevent sedimentation and measured with a SpectraMax M5 (Molecular Devices) every 30 minutes. Each strain/medium combination was grown in 3 separate plates with 2 technical replicates per plate.

For RNA isolation 10 mL cultures were grown in 15 mL falcon tubes at 37 °C with 5% CO_2_. All experiments were carried out using a single batch of THY and APS. Based on the growth curves we identified a timepoint where all strains were in the (mid) exponential growth phase. To account for different growth in falcon tubes compared to 96-well plates, we reduced the incubation time and confirmed that the culture was in the correct growth phase by measuring OD_600_ before RNA isolation. Cultures in APS were sampled at 95 minutes and THY cultures at 125 minutes. The APS and THY cultures were started at different times to enable RNA isolation at the same time. Cultures were pelleted by centrifugation and resuspended in QIAzol Lysis Reagent (Qiagen). After bead beating twice for 40 s with 0.1 mm silica beads (MP biomedicals) RNA was isolated using the miRNeasy kit (Qiagen). Trace DNA was digested using the DNase Max Kit (Qiagen). RNA quantity and integrity was confirmed with nanodrop (ThermoFisher) and TapeStation (Agilent). Library preparation was done with the Illumina Stranded Total RNA Prep kit. rRNA was enzymatically depleted and the remaining RNA fragmented and translated to cDNA. The libraries were sequenced with 150 bp paired-end sequencing on the Illumina NovaSeq 6000 system with Illumina NovaSeq 6000 SP reagent kit at iGenSeq (Institut du Cerveau, France).

### Bioinformatics

The strain genomes were annotated with Prokka v1.14.5 [23]. We further identified antimicrobial resistance genes with Resfinder v4.1 [24] and biosynthetic gene clusters with antiSMASH v5.1.2 [25] and BiG-SCAPE v1.1.2 [26]. We used clinker [27] for gene cluster comparison. To compare the transcriptome between strains we grouped homologous and paralogous genes using Orthofinder v2.3.12 [28, 29], and the gene expression for paralogous genes was summed. This may in some cases result in the grouping of genes with different function. The “orthogroups” found by Orthofinder were given static names from the RefSeq locus tags of *S. suis* reference strains P1/7 (GCF_000091905.1) and D12 (GCF_000231905.1). GO terms for each Orthogroup were found with InterProScan v5.54-87.0 [30], and quantitative analysis of GO term expression was done on the whole transcriptome, including accessory genes. The RNA sequencing data was adapter and quality trimmed with Trimmomatic v0.39 [31] before being mapped to the genome of the individual strain with bowtie2 [32]. FeatureCounts 2.0.1 [33] with default settings was used to count the number of reads mapping to each gene. Further analysis was carried out with R v4.1.3 [34].

### Analysis

Combining the transcriptomes of different strains into a single analysis requires additional normalization compared to single-strain transcriptomic studies. In addition to variable sequencing depth per sample, strains vary in genome size and presence/absence and length of genes. We opted to use TPM read count normalization to facilitate comparison between strains. We identified differentially expressed genes using Wilcoxon Rank Sum Test with FDR correction. Overall transcriptome difference was calculated separately using the whole genome transcriptome (expression of accessory genes absent in strain set to 0) and the core genome transcriptome (using only core genes shared by all strains). To quantify and visualise overall transcriptome difference we used R package vegan [35] Bray-Curtis dissimilarity, principal component analysis (PCA), and redundancy analysis (RDA). To determine if the transcriptome conservation differed significantly between groups, we used estimated marginal means on linear models with R function emmeans [36].

## Results and discussion

### Dataset description

We sequenced the transcriptome of a genetically diverse set of 21 *S. suis* strains (Figure 1A) in the mid exponential growth phase in Todd Hewitt Yeast Broth (THY) and active porcine serum (APS). Overall, commensal and pathogenic strains grew equally well in both media (Figure S1). All samples were successfully sequenced with a minimum of 20 million 150 bp paired-end reads (1114-1902x coverage). All samples were free from contamination, with >99.8% reads mapping to the corresponding strain genome. One sample, S11 grown in THY, had excessively high expression of the arginine deiminase system (ADS, Figure S2). ADS is previously described as vital for *S. suis* growth in acidic medium [37, 38], and the high ADS expression may be linked to *in vitro* culturing and have limited *in vivo* relevance. Considering the excessive impact of this single gene cluster on the total transcriptome we excluded the sample from the main analysis. Table S2 shows TPM values per orthogroup for all sequenced samples.

**Figure 1:**
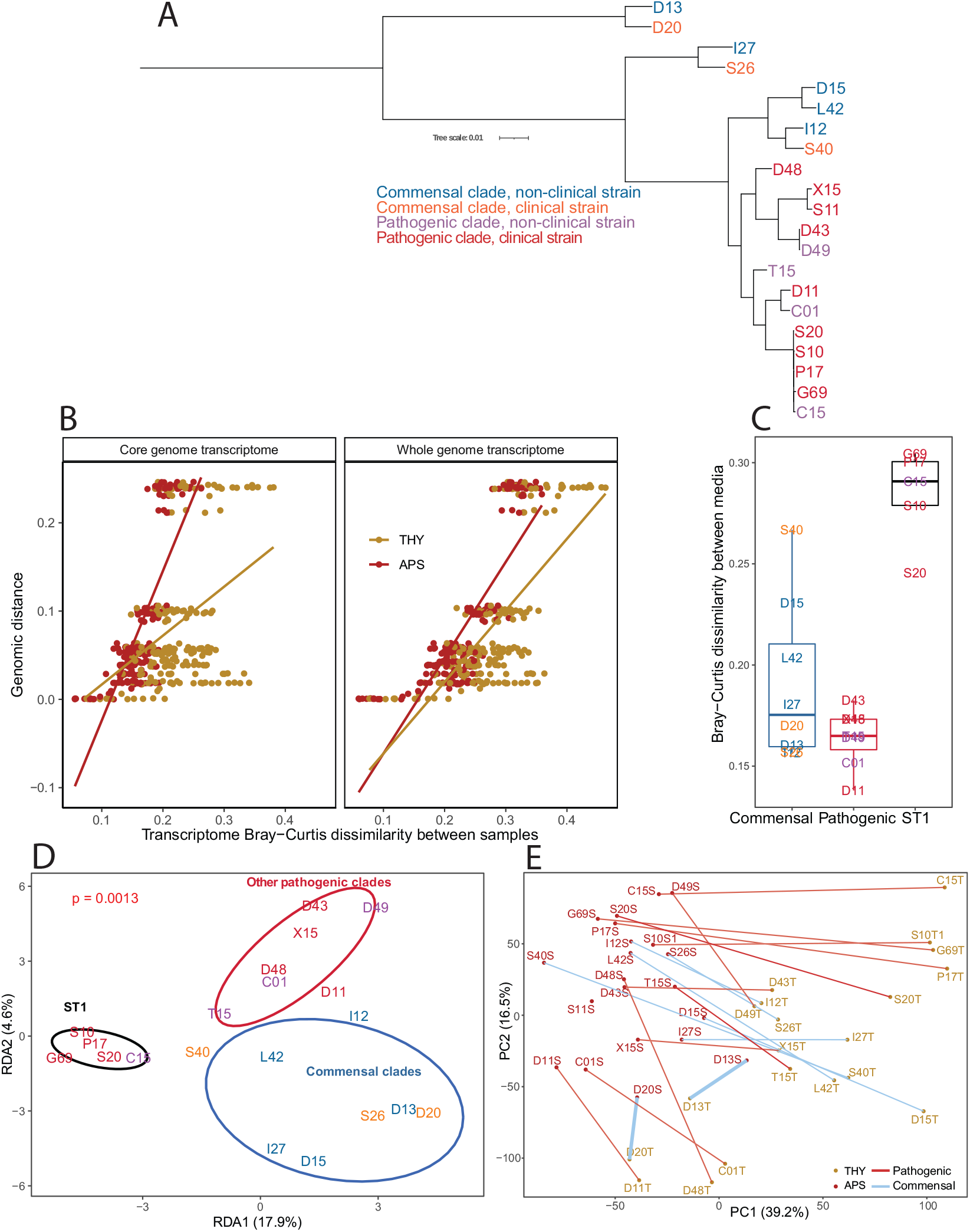
Strain phylogeny and overall transcriptome variation. **A)** Core genome phylogenetic tree of the 21 strains included in the study, constructed by Orthofinder and STAG with default settings and mid-point rooted. While ST1 pathogenic strains such as P1/7 and S10 are closely related, commensal clades have high genomic diversity. **B)** Transcriptome Bray-Curtis dissimilarity between strains correlates with genome phylogenetic distance. The transcriptome is more conserved in APS than in THY. Each point represents a pairwise comparison. Strains were only compared within the same medium. **C)** Boxplot of core genome transcriptome Bray-Curtis dissimilarity between samples from each strain in the two media. **D)** Redundancy analysis (RDA) on log2(fold-change) on the core genome transcriptome constrained by [ST1 vs other pathogenic clades vs commensal clades] showed that ST1 clade strains had a distinct transcriptome change between media. Eclipses represent 75% confidence level. **E)** Principal component analysis (PCA) showing core genome transcriptome separation between samples grown in THY and APS. The pairs of samples were well separated on PC1 in all strains except for strain D13 and D20. Each label (coloured by medium) is one sample, and the two samples of each strain are joined by lines coloured by clade type.

Comparison of S10 triplicate samples showed that our methods were highly reproducible, with a maximum pairwise Bray-Curtis dissimilarity of 0.1 between replicates. Distinct but very closely related strains within sequence type 1 (ST1) had similar pairwise dissimilarities, but this was expected as their genomes are virtually identical. It does, however, show that small differences in gene expression between strains should be interpreted with care. For the remaining analysis only a single S10 sample from each medium was included.

### Serum opacification

Three commensal strains increased the OD_600_ rapidly and without lag phase when grown in APS, reaching far higher OD_600_ values than the other strains (Figure S1). This was not due to an increase of bacterial biomass, but opacification of the supernatant. Formation of large lipid particles by the protein opacification factor of serum (*ofs*, SSU_RS07445) has previously been described in *S. suis,* and the gene is known to occur in different variants with and without opacifying function [39, 40]. The three strains with strong opacification activity had high expression of *ofs* variants diverged from those previously described (Figure S3). While the shorter *ofs* type-1 [40] variant found in ST1 is expressed at low levels and has limited opacification capacity, it is conserved in pathogenic clades and may have a function related to binding to host cells [39, 41].

### Conservation of core genome expression in serum

Commensal clades are diverse, with long (core- and whole genome-based phylogeny) branch lengths between strains, while pathogenic clades, in particular the commonly researched ST1 clade, consist of closely related strains [6]. Our strain selection included both ST1 strains and a wider range of strains from other pathogenic and commensal clades. Strains D13 and D20 form an outgroup to the other strains in our dataset (Figure 1A). The phylogenetic status of this “divergent” outgroup clade is debated, as it can also be considered a separate species (see closely related strains in “clade 2”, Baig et. al. 2015) [42]. Despite large genomic differences (less than 86% ANI) to other included strains, their transcriptome was conserved in APS (measured in Bray-Curtis dissimilarity, Figure 1B). Overall, the *S. suis* transcriptome was significantly more conserved in APS compared to THY, and in the expression of the core genome compared to the whole genome transcriptome (estimated marginal means, p < 0.01). Despite the large genomic differences between outgroup D13+D20 and the remaining strains, pairwise transcriptome Bray-Curtis dissimilarities to the other strains was overlapping with that between strains from different pathogenic clades. The 13 GO terms most expressed in THY were all more expressed in APS (Figure S4). This suggests that the conserved APS transcriptome reflects upregulation of core physiological functions related to growth and cell division. This may be relevant in facilitating exponential growth during host invasion. See table S3 for differences in expression of all GO terms and genes between clades and media.

### Large transcriptome differences between media in ST1 strains

Transcriptome difference between the two media was largest for ST1 strains (Figure 1C). Redundancy analysis (RDA) on log2fold change of gene expression between THY and APS cultures constrained by clade type (ST1 vs other pathogenic clades vs commensal clades) also showed that regulation of gene expression in ST1 strains was distinct, while other pathogenic clades and commensals were more similar (Figure 1D). The transcriptome of ST1 strains was, however, not strongly divergent from other clades in either medium. PCA on all samples (Figure 1E) showed that the pairs of samples from each strain separated in similar directions on the PC1 and PC2 axis, except for the two outgroup commensal strains D13 and D20. ST1 strains showed stronger regulation of several genes. This included downregulation of SSU_RS07200 (SprT-like protein) and upregulation of SSU_RS02105 (cysteine synthase) in THY (Figure S5). ST1 and other pathogenic *S. suis* are thought to regulate gene expression related to virulence factors via a phase variable type I DNA methyltransferase system [9, 43]. However, this system is unlikely to explain the differences seen in the present study as the THY and APS cultures grew separately for only a few generations. This is unlikely to be sufficient time for selection to drive change across the experimental population. We did not observe differences in growth rates of the strains, indicating that selection pressure for either phase variant was limited, and that phase variation did not occur. Moreover, non-ST1 strains with the same phase variation system had small transcriptome differences between media. These results suggest that ST1 strains may have additional undescribed mechanisms enabling strong regulation of gene expression in different environmental conditions.

### High *cps* expression in serotype 2 strains

The *S. suis* polysaccharide capsule (CPS) exists in many variants (serotypes). Serotype 2, a capsule type with terminal sialic acid, is the most studied serotype due to its association with high (zoonotic) virulence [44–46]. In this study all ST1 strains were serotype 2, in addition to strains T15 and D43+D49. Genes involved in capsule production (cps) were upregulated in APS in most strains (Figure 2A). *Cps* expression was high and strongly upregulated in APS in serotype 2 strains compared to other serotypes, although serotype 2 strains D43 and D49 appeared to have divergent regulation. D49 downregulated *cps* expression less, and D43 had higher *cps* expression in THY, opposite of other serotype 2 strains. Non-typeable strains D13 and D20 had the lowest *cps* expression, but apart from these and serotype 2 strains, *cps* expression overlapped between commensal (serotype 4, 16, 19, 30, and 31) and pathogenic clades (serotype 7, 8, and 9). The pathogenic clades had higher expression levels than the commensals, but this was only significant in APS (p = 0.03) and not in THY (p = 0.15).

**Figure 2:**
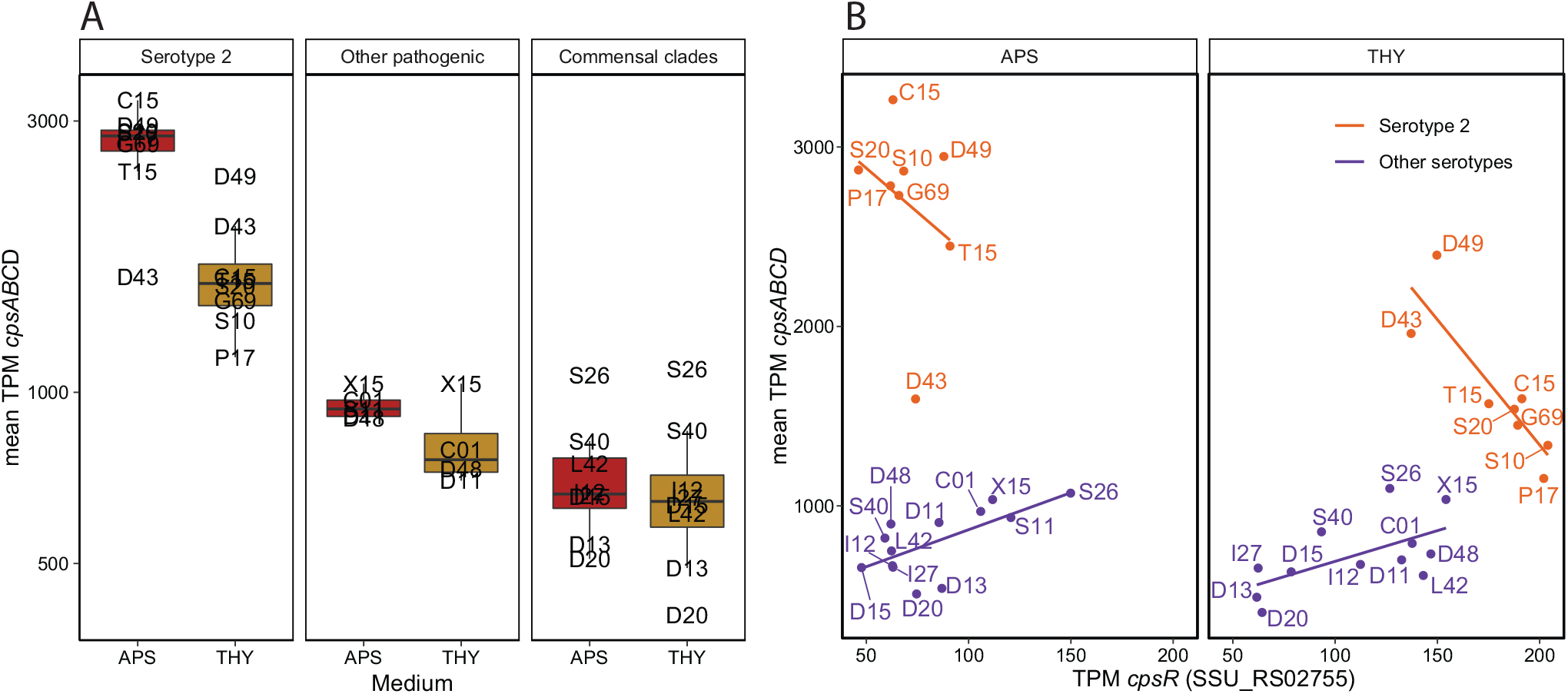
Capsule gene cluster expression. **A)** Mean expression of *cpsABCD* (SSU_RS02765-SSU_RS02780), the initial 4 genes in the cps gene cluster which are shared by all strains. Grouped by serotype 2 (all from pathogenic clades), other pathogenic clade strains, and commensal clade strains. **B)** Scatterplot with regression lines comparing the mean expression of *cpsABCD* with the *cpsR* (SSU_RS02755) regulator. Only the serotype 2 strains had a negative correlation between *cpsR* and cps expression, although serotype 2 strains D43 and D49 also appeared to be regulated differently as *cpsABCD* expression was similar in the two media.

Streptococcal *cps* expression has been linked to negative regulation by a protein (SSU_RS02755) variably named as *cpsR*/*gntR*/*orf2Y* [47–49]. In *S. pneumoniae, cpsR* has been shown to interact with the *cps* promoter dependent on glucose concentration, negatively controlling *cps* expression and CPS production [48]. We found a negative correlation between *cpsR* and *cps* expression only in serotype 2 strains (Figure 2B). This, and the lack of *cps* regulation in the serotype 2 strains D43 and D49, may be due to variation in *cpsR*, which occurs as a distinct variant in all serotype 2 strains except D43 and D49. Although the D43 and D49 *cpsR* sequences contain arginine in position 17 characteristic for serotype 2 strains, all other sequence variation specific to this serotype is absent (Figure S6, S7). It is not clear how different combinations of serotypes and *cps* regulation may be beneficial in different niches.

### High expression of the agmatine deiminase system in serum in commensal strains

As one may expect, many of the genes and pathways most differentially expressed between THY and APS related to acquisition and metabolism of nutrients that are differentially present in the media (Figure 3A, Table S3). Metal ion acquisition was upregulated in APS while trehalose import and metabolism were highly expressed in some strains in THY. All strains had high expression of the *lac* operon (downstream from SSU_RS04590) in THY, but this was strongly downregulated in APS, except in outgroup strain D13. This strain expressed the *lac* operon constitutively, possibly due to having an unusual, truncated, copy of encoding repressor *lacR.* It did, however, also carry a *lac* gene cluster copy more similar to other strains (Figure S8). Lactose metabolism is likely to be important for *S. suis* during commensal colonisation of pre-weaning piglets, and the fitness cost of constitutive *lac* operon expression is likely limited (D13 was collected from a piglet approximately 42 days old and thus appears able to persist post-weaning).

**Figure 3:**
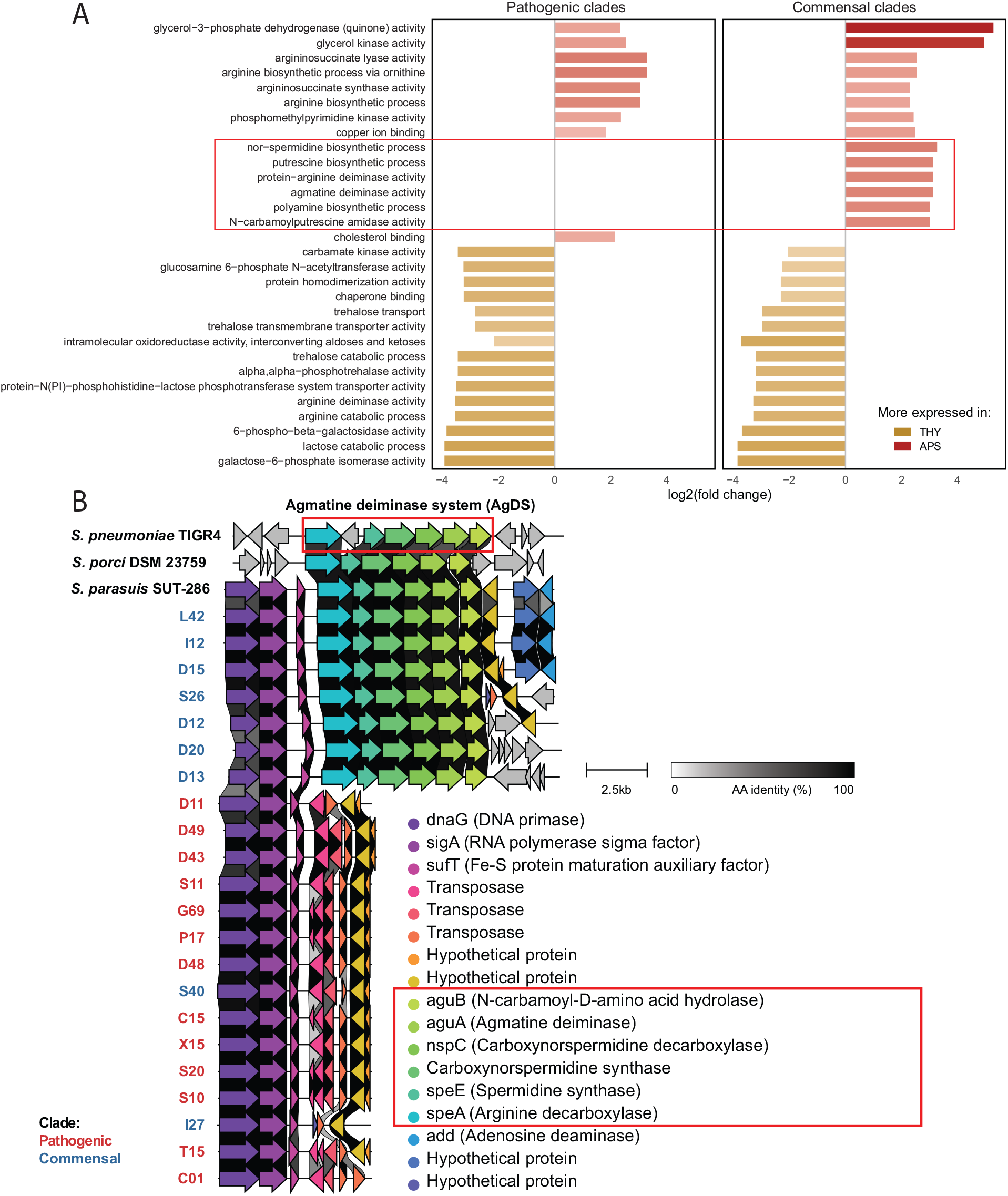
The accessory AgDS system has a large influence on transcriptome differences between THY and APS in commensals. **A)** The 30 GO terms with the largest log2fold changes between THY and APS cultures (average across all strains, GO terms with <10 mean TPM excluded). Several GO terms highly expressed in APS occurred only in commensal clades due to presence/absence of the agmatine deiminase system, suggesting that this gene cluster may be important for commensal *S. suis.* GO terms associated with the agmatine deiminase system highlighted in red. **B)** The AgDS gene cluster is found only in commensal clade strains and has high similarity to the AgDS of S. *pneumoniae, S. porci,* and S. *parasuis.*

One of the largest overall differences between growth in THY and APS was in amino acid metabolism. The genomes of 6 of 8 commensal strains, but no pathogens, encode an agmatine deiminase system (AgDS, SSUD12_RS06980-SSUD12_RS07005). In the strains in which it was present, AgDS was highly expressed and upregulated in APS compared to in THY. This gene cluster has not previously been described in *S. suis* but is known from *S. pneumoniae* [50] and *S. mutans* [51–53]. The *S. suis* AgDS share high similarity to the AgDS of *S. pneumoniae* reference strain TIGR4 (SP_RS04525-SP_RS04560), *S. porci,* and *S. parasuis* (Figure 3B). The *S. mutans* AgDS is diverged, with several indels and less than 50% amino acid residue identity to the *S. suis* agmatine deiminase gene. In *S. suis* the AgDS appear to be linked to a set of unrelated genes including DNA primase (Figure 3B), and in some strains these make up a 11.5 kb genomic island flanked by transposases.

The arginine deiminase system (ADS/*arcABC* operon, SSU_RS03045-SSU_RS03060) was found in all strains and in contrast to AgDS it was more expressed in THY than in APS. Agmatine is formed upon decarboxylation of arginine, and agmatine and arginine metabolic pathways show considerable overlap. Both the arginine and agmatine deiminase system produce ammonia, increasing intracellular pH and tolerance to low pH. High ADS and AgDS expression may be linked to acidification of the medium during growth. ADS has been shown to be important for *S. suis* survival in acidic medium [37, 54]. AgDS is thought to be relevant to acidic stress in *S. mutans* [53], but does not appear to increase pH tolerance in *S. pneumoniae* [50]. It is possible that AgDS activity provides a competitive advantage to commensal *S. suis* by increasing intracellular pH. Agmatine released by competing taxa inhibits the growth of *S. mutans* [51], and a similar mechanism may apply to *S. suis.* Considering its variable presence, it may be relevant to both inter- and intra-species competition.

### High expression of Mn^2+^ binding lipoprotein *troA* in pathogens

Transition metal ion homeostasis is vital to bacteria. While commensals sequester metal ions in competition with other microbes, pathogens must overcome host metal ion chelation during infection. Some lactic acid bacteria, including *S. suis* and S. *pneumoniae,* utilize a larger proportion of manganese relative to iron than most bacteria, and this may provide them with a competitive advantage in some niches [55–58]. We found that genes putatively involved in both iron (SSU_RS03155-SSU_RS03170) and manganese (SSU_RS09395-SSU_RS09415) scavenging were more expressed in APS than THY, although the manganese import gene cluster was 5 times more expressed than the iron import gene cluster. *TroA,* a putative scavenger protein for the *troBCD* ABC transport system [58–60], was also 3-4 times more expressed than the *troBCD* ABC transporter genes in the gene cluster (Figure 4A). This indicates that expression of this gene is regulated separately.

**Figure 4:**
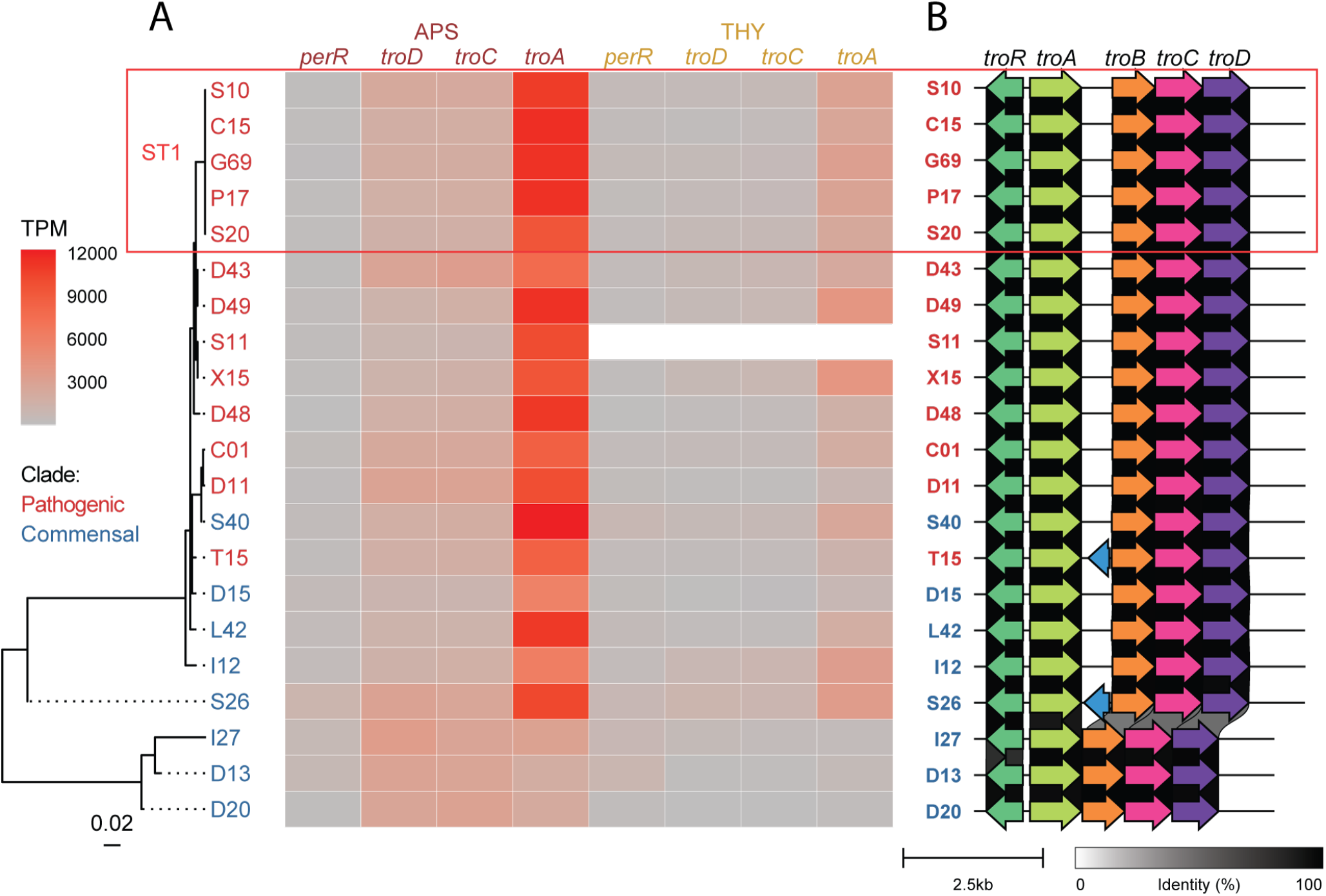
Variation in sequence and expression level of manganese import gene cluster *troABCD* and its regulators. **A)** Regulator *perR* maximum-likelihood tree with heatmap of gene expression TPM values. Mn^2+^ binding lipoprotein *troA* was more expressed than *troBCD* and more expressed in APS compared to THY. Some commensal strains with diverged *perR* and *troRABCD* copies had reduced *troA* expression. **B)** In addition to sequence variation to the other strains, strains I27, D13, and D20 shared a truncated *troABCD* gene cluster which lacked SSU_RS09410, a putative membrane protein variably annotated as a pseudogene (shown in blue for T15 and S26) or unannotated by Prokka in different strains, likely due to its short length and truncation.

*TroA* expression varied greatly between strains (232-4165 TPM in THY and 1899-12228 TPM in APS). Some commensals had reduced t*roA* expression compared to ST1 strains despite having similar or higher *troBCD* expression (Figure 4A). This may be due to sequence variation of previously described regulators and the gene cluster itself. *S. suis troABCD* expression has been reported to be repressed by *dtxR* family metalloregulator *troR* (SSU_RS09420) and fur family regulator *perR* (SSU_RS01575) depending on metal ion concentrations and oxidative stress [15, 60–62]. Additionally, *mntE* (SSU_RS05010) has been identified as a manganese efflux system [63]. Both *troR* and *perR* may contribute to *troABCD* upregulation in APS compared to THY. Variation in *troA* expression levels between strains grown in the same medium are unlikely to be caused by *troR*, because strains with identical variants varied greatly in expression. *PerR* and *troA* itself also had sequence variation, and the strains with the lowest *troA* expression had a truncated gene cluster structure (Figure 4B, Figure S9). A region between *troA* and *troB* contains a putative oligopeptide transporter pseudogene in most strains, but this was deleted in the strains with the lowest *troA* expression, D13, D20, and I27 (Figure 4B). The initial part of this region has higher expression than *troA,* and its conservation across most of the included *S. suis* clades suggests that it may have a function irrespective of (the length of) the longest predicted open reading frame.

## Conclusions

We found that *S. suis* strains with large genomic divergence have unexpectedly conserved transcriptomes when grown in APS. More variation was observed in THY, and this should be considered when selecting growth medium for *in vitro* assays. Despite overall conservation, the transcriptome of the strains varied in key functions, including well described regulatory mechanisms. In most clades the manganese import and capsule gene clusters may be regulated differently than described for ST1 strains. In general, ST1 strains displayed larger changes in gene expression between THY and APS cultures compared to other clades, and these differences in gene expression may help them rapidly adapt to environmental changes, most notably when changing between upper respiratory tract colonization and host invasion. The gene cluster encoding production of the capsule, which is key in avoiding the host complement attack complex, was among the genes most strongly upregulated and highest expressed in ST1 strains.

## Data summary

The genome assemblies generated in this study are available under BioProject PRJNA855487. The BioSample accession number of each strain is listed in Table S1. RNA-sequencing data is available under BioProject accession number PRJNA863843 (not released at preprint publication).

## Acknowledgements

We thank Maria Laura Ferrando and Isabela Fernandes de Oliveira for providing strains and Illumina sequencing data.

This research was financially supported by EU Horizon 2020 Program Grant Agreement 727966), funded under H2020-EU.3.2.1.1. SF is a PhD student funded by the Netherlands Centre for One Health.

