## Supplementary figures for "Transcriptomics in serum and culture medium reveal shared and differential gene regulation in pathogenic and commensal *Streptococcus suis*"

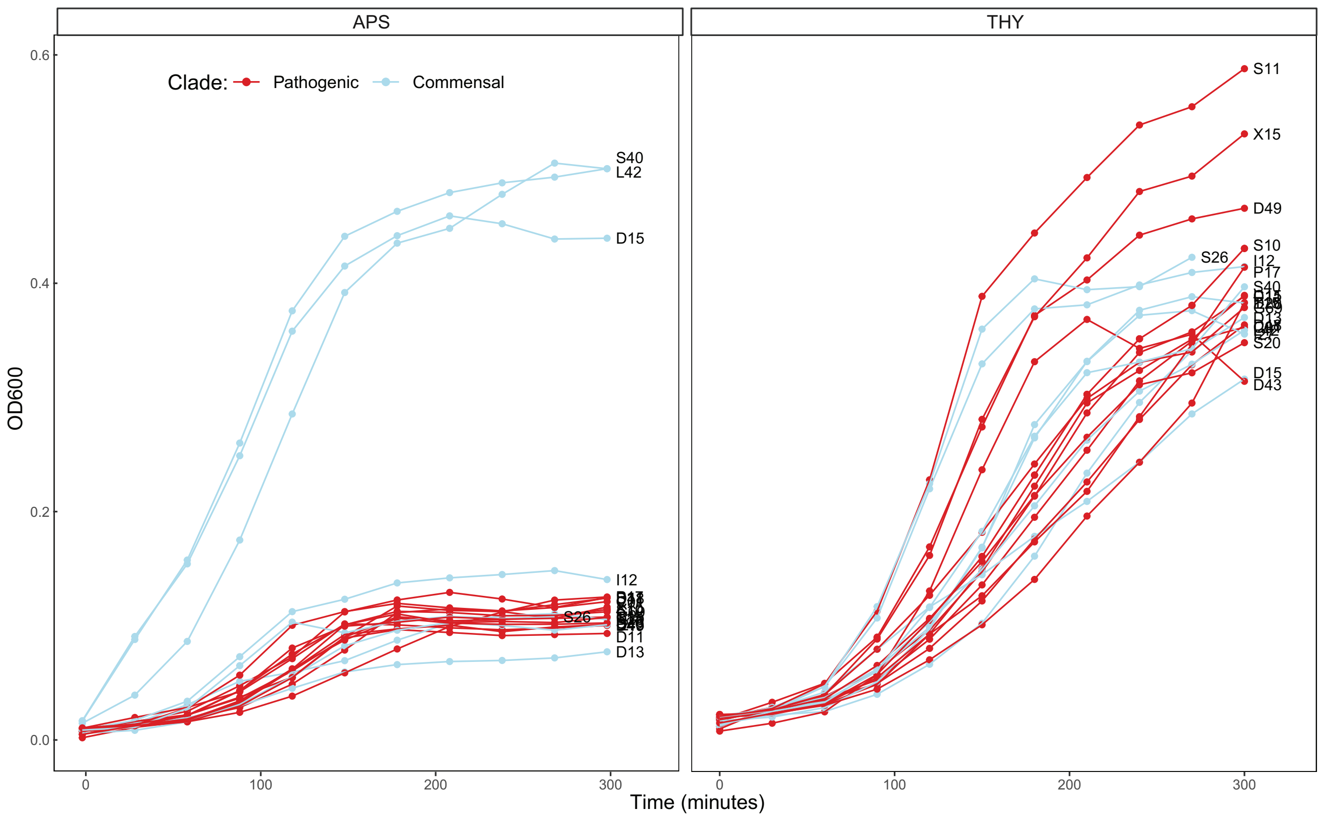


**Figure S1:** Growth curves. The strains were grown in 96-well plates with 200 µL total volume per well and incubated at 37 °C with 5% $CO_2$. Each point represents the mean value of the strain sampled in minimum 3 separate plates with 2 technical replicates per plate.

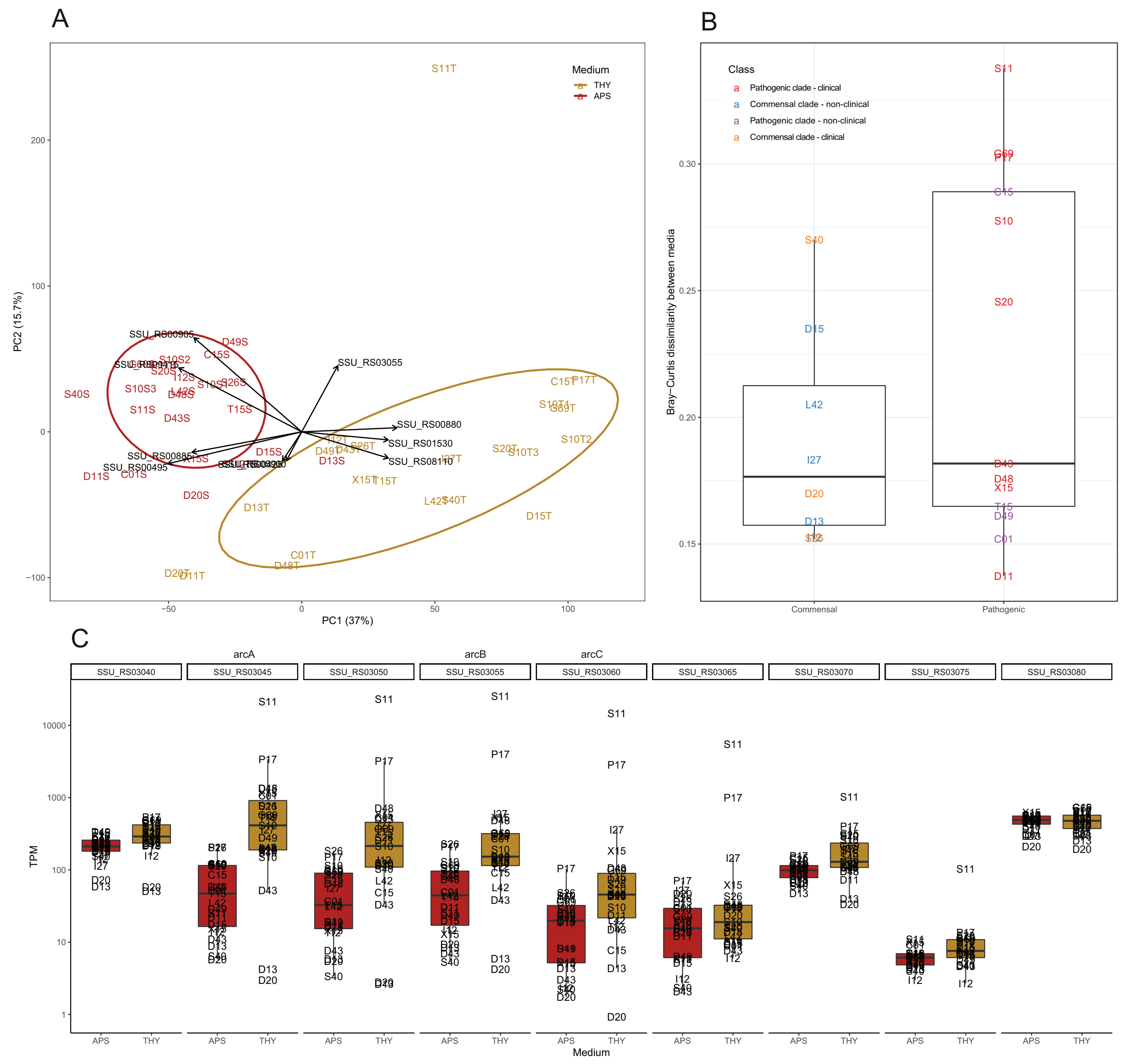


**Figure S2:** High ADS expression in S11 grown in THY. **A)** PCA on the core genome transcriptome of all strains shows sample S11T (strain S11 grown in THY) as an outlier. Each label represents a single sample, eclipses represent 75% confidence level, arrows show genes driving the separation of samples; samples in the direction the arrow is pointing have higher expression of the gene. **B)** Strain S11 also had the largest transcriptome difference between THY and APS cultures of the same strain (core genome transcriptome Bray-Curtis dissimilarity). **C)** Boxplot of expression (shown as log10(TPM) of *arcABC* and adjacent genes.

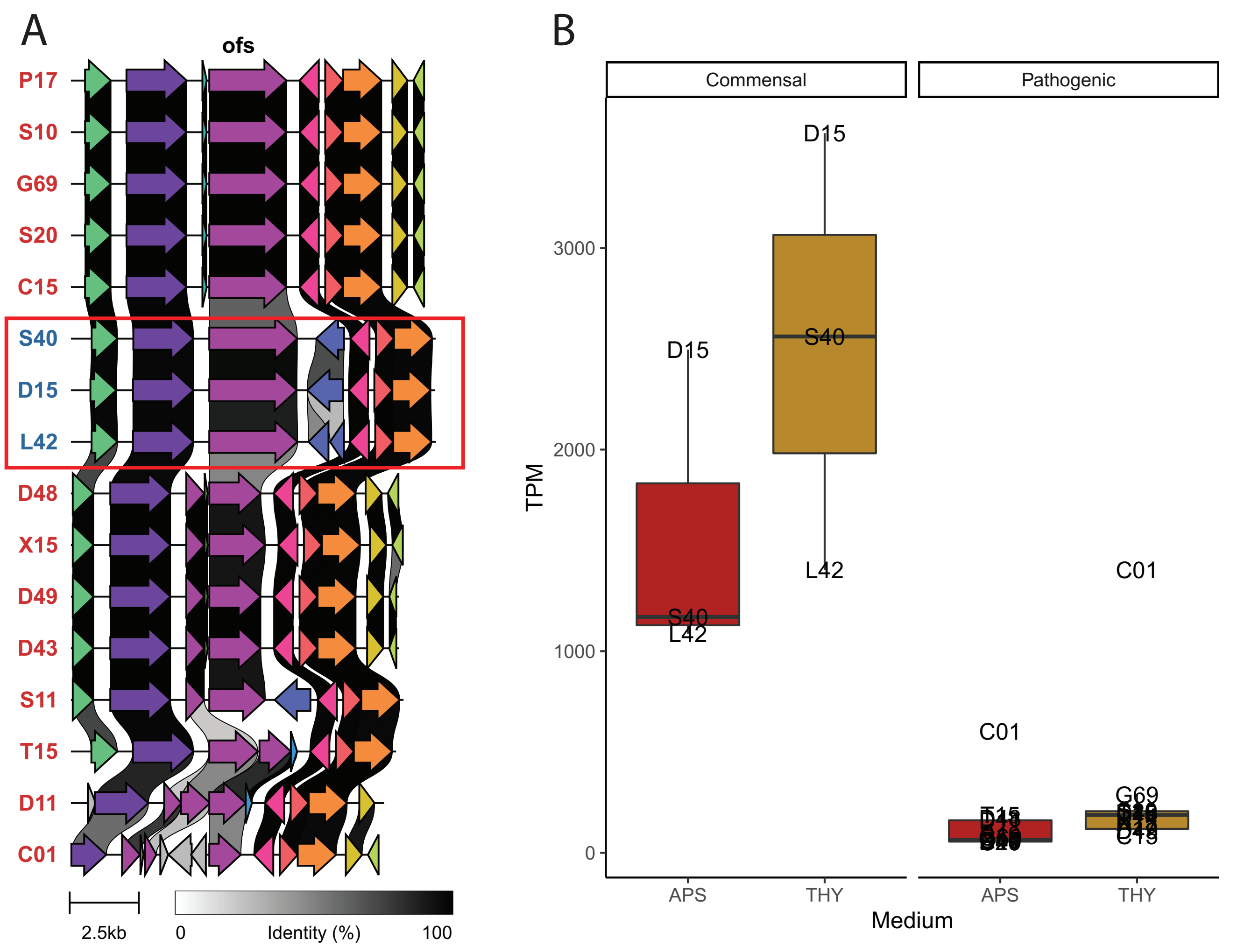


**Figure S3:** Opacification factor of serum (*ofs*). **A)** *ofs* occurs in different (suggested pseudogene) variants, but in similar genomic context. The 3 strains with high opaficification activity in serum (red box) share a similar variant (although there is substantial nucleotide level variation between these 3 strains). **B)** The 3 commensals with *ofs* had high expression of the gene. The other commensal strains did not have any detected *ofs* homologue.

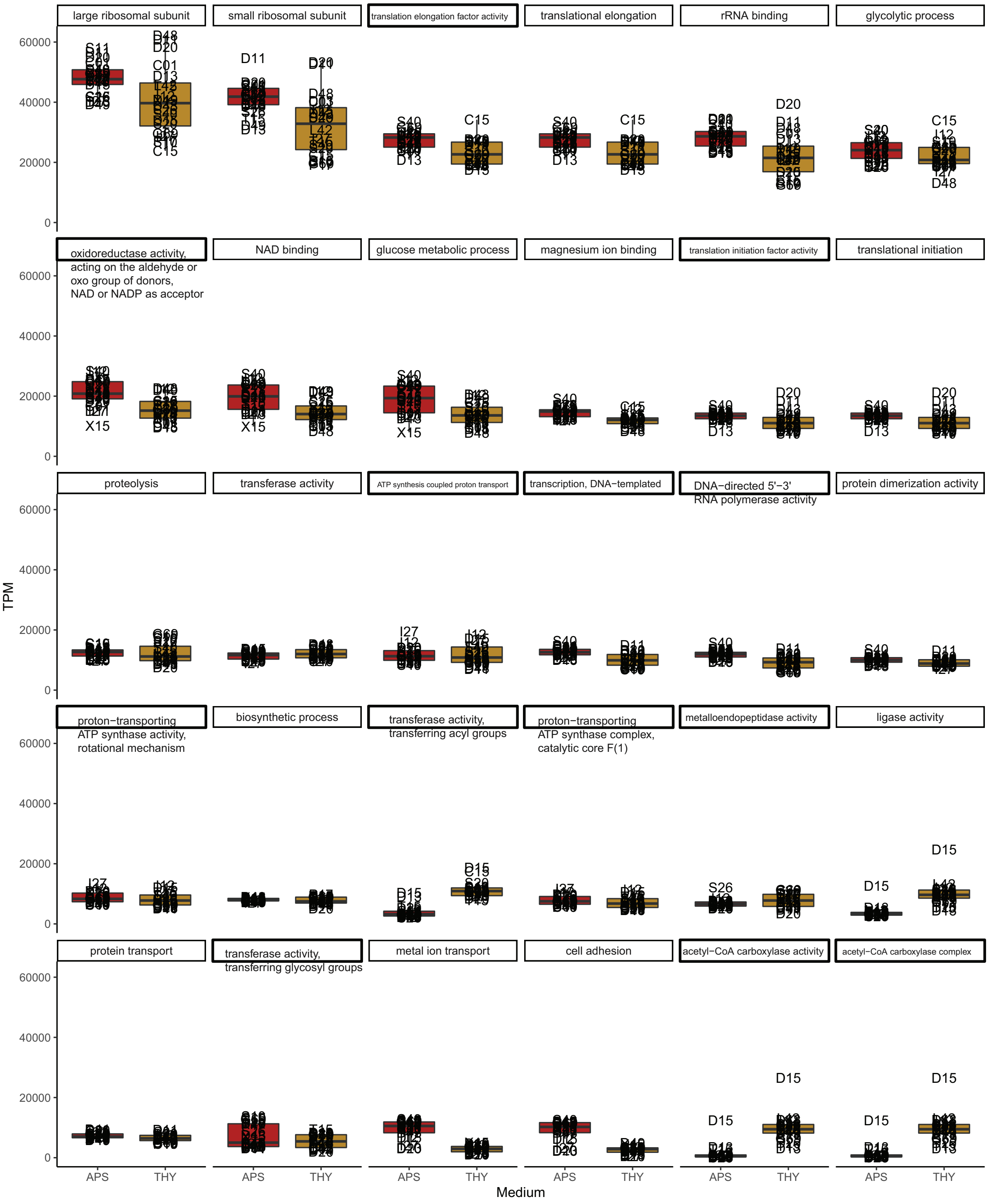


**Figure S4:** Boxplots showing expression of the overall most expressed GO terms in APS vs THY.

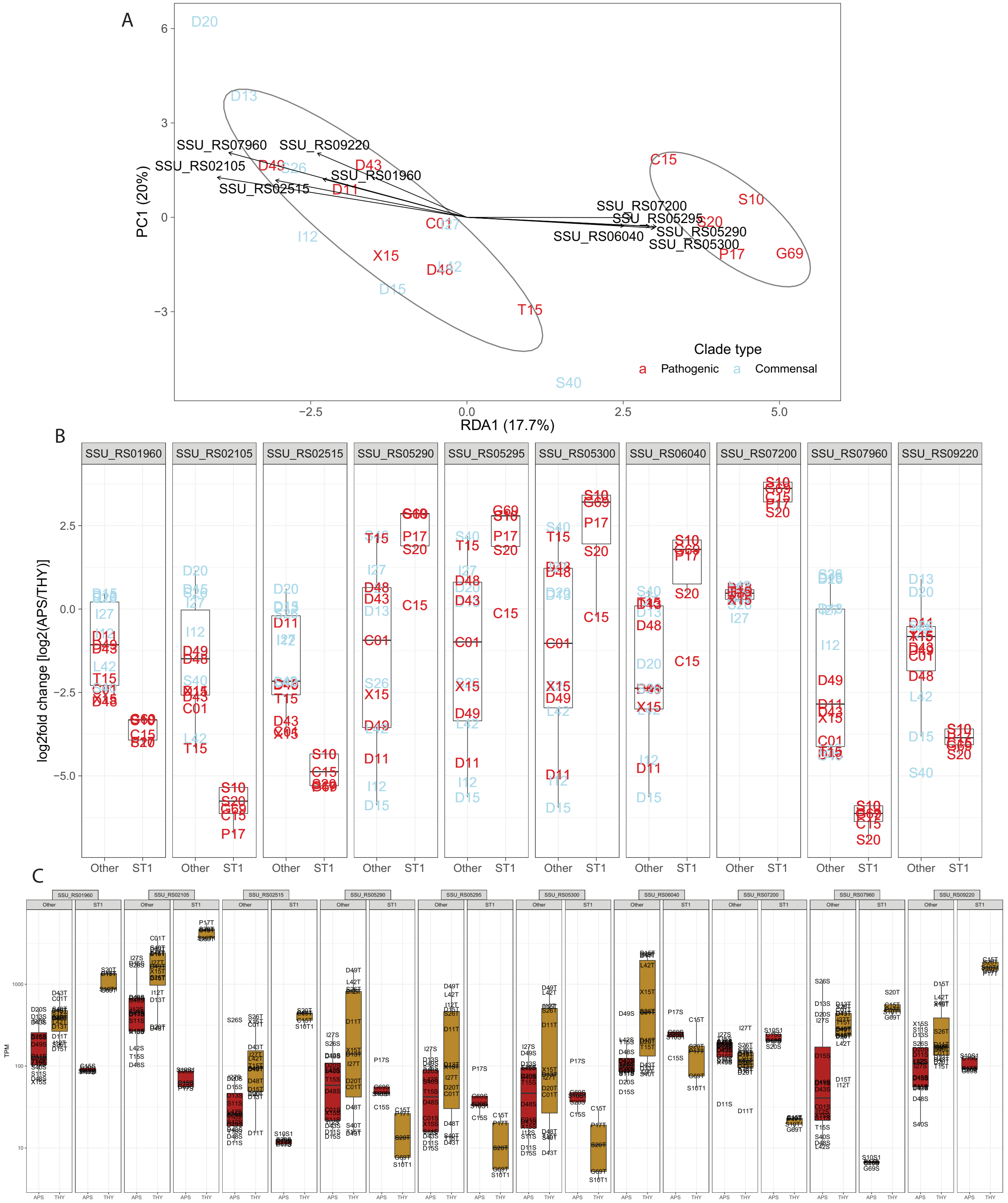


**Figure S5:** Genes strongly regulated between media in sequence type 1 strains. **A)** RDA on log2fold gene expression change constrained by ST1 clade strains vs all other strains. Each label represents a single sample, eclipses represent 75% confidence level, arrows show genes driving the separation of samples; samples in the direction the arrow is pointing have higher expression of the gene. **B)** Boxplot of the log2fold change of the 10 strongest drivers (genes) of the RDA1 axis. **C)** Boxplot of the TPM expression of the same 10 genes in APS and THY.

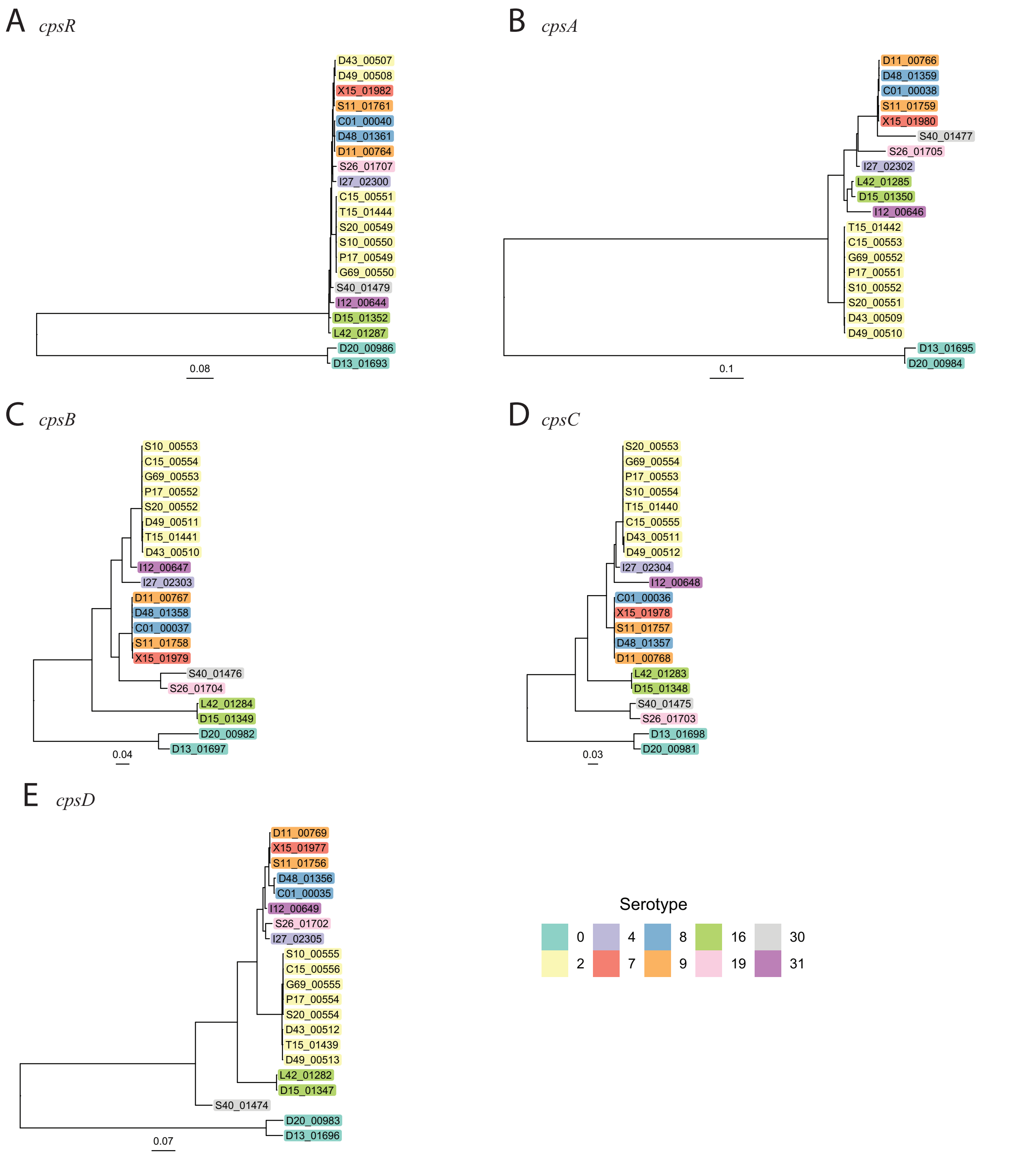


**Figure S6:** Maximum likelihood trees of *cps* genes (DNA). **A)** SSU_RS02755, *cpsR/gntR/orf2Y* **B)** SSU_RS02765, *cpsA* **C)** SSU_RS02770, *cpsB* **D)** SSU_RS02775, cpsC **E)** SSU_RS02780, *cpsD*. Tip labels: [Strain]_[gene number]. Except from outliers D13 and D20, *cpsR* is more conserved than *cpsABCD*. Diverged *cpsR* copies in strains D43 and D49 may underlie different *cps* regulation compared to other serotype 2 strains.

```
C15       ..MKKYVDTGHRKRGKSRRYHCSKDTVKAMKYNKYAVKSGYYDRDDHTVNTDYDRCKSGRNYNYYHGASSVSMDYHVYTKKDVTAGSAYT   88
G69       ..MKKYVDTGHRKRGKSRRYHCSKDTVKAMKYNKYAVKSGYYDRDDHTVNTDYDRCKSGRNYNYYHGASSVSMDYHVYTKKDVTAGSAYT   88
P17       ..MKKYVDTGHRKRGKSRRYHCSKDTVKAMKYNKYAVKSGYYDRDDHTVNTDYDRCKSGRNYNYYHGASSVSMDYHVYTKKDVTAGSAYT   88
S10       ..MKKYVDTGHRKRGKSRRYHCSKDTVKAMKYNKYAVKSGYYDRDDHTVNTDYDRCKSGRNYNYYHGASSVSMDYHVYTKKDVTAGSAYT   88
S20       ..MKKYVDTGHRKRGKSRRYHCSKDTVKAMKYNKYAVKSGYYDRDDHTVNTDYDRCKSGRNYNYYHGASSVSMDYHVYTKKDVTAGSAYT   88
T15       ..MKKYVDTGHRKRGKSRRYHCSKDTVKAMKYNKYAVKSGYYDRDDHTVNTDYDRCKSGRNYNYYHGASSVSMDYHVYTKKDVTAGSAYT   88
S40       ..MKKYVDTGHRKRGKSRSYHCSKDTVKAMKYNKYAVKSGYYDRDDHTVNADYDRCNSGRNYNYYHGASSVSMDYHVYTKKDVTAGSAYT   88
I12       ..MKKYVDTGHRKRGKSRSYHCSKDTVKAMKYNKYAVKSGYYDRDD.TVNADYDRCNSGRNYNYYHGASSVSMDYHVYTKKDVTAGSAYT   87
D15       ..MKKYVDTGHRKRGKSRSYHCSKDTVKAMKYNKYAVKSGYYDRDDHTVNADYDRCNSGRNYNYYHGASSVSMDYHVYTKKDVTAGSAYT   88
L42       ..MKKYVDTGHRKRGKSRSYHCSKDTVKAMKYNKYAVKSGYYDRDDHTVNADYDRCNSGRNYNYYHGASSVSMDYHVYTKKDVTAGSAYT   88
S26       ..MKKYVDTGHRKRGKSRSYHCSKDTVKAMKYNKYAVKSGYYDRDDHTVNADYDRCNSGRNYNYYHGASSVSMDYHVYTKKDVTAGSAYT   88
D43       ..MKKYVDTGHRKRGKSRRYHCSKDTVKAMKYNKYAVKSGYYDRDDHTVNADYDRCNSGRNYNYYHGASSVSMDYHVYTKKDVTAGSAYT   88
D49       ..MKKYVDTGHRKRGKSRRYHCSKDTVKAMKYNKYAVKSGYYDRDDHTVNADYDRCNSGRNYNYYHGASSVSMDYHVYTKKDVTAGSAYT   88
X15       ..MKKYVDTGHRKRGKSRSYHCSKDTVKAMKYNKYAVKSGYYDRDDHTVNADYDRCNSGRNYNYYHGASSVSMDYHVYTKKDVTAGSAYT   88
I27       ..MKKYVDTGHRKRGKSRSYHCSKDTVKAMKYNKYAVKSGYYDRDDHTVNADYDRCNSGRNYNYYHGASSVSMDYHVYTKKDVTAGSAYT   88
C01       ..MKKYVDTGHRKRGKSRSYHCSKDTVKAMKYNKYAVKSGYYDRDDHTVNADYDRCNSGRNYNYYHGASSVSMDYHVYTKKDVTAGSAYT   88
D11       ..MKKYVDTGHRKRGKSRSYHCSKDTVKAMKYNKYAVKSGYYDRDDHTVNADYDRCNSGRNYNYYHGASSVSMDYHVYTKKDVTAGSAYT   88
D48       ..MKKYVDTGHRKRGKSRSYHCSKDTVKAMKYNKYAVKSGYYDRDDHTVNADYDRCNSGRNYNYYHGASSVSMDYHVYTKKDVTAGSAYT   88
S11       ..MKKYVDTGHRKRGKSRSYHCSKDTVKAMKYNKYAVKSGYYDRDDHTVNADYDRCNSGRNYNYYHGASSVSMDYHVYTKKDVTAGSAYT   88
D13       ...MKYKDDGRDYKRGKSRSYCSKDTVKAMKYNKYAVKSGYYVDDR.DDDTDYDRCHSGRNYNYYHAGTTSVMDYHVYSRKDVTAGSAYS   86
D20       MSKYRDVDDGRDYKRGKSRSYCSKDTVKAMKYNKYAVKSGYYVDDH.DDDTDYDRCHSGRNYNYYHAGTTSVMDYHVYSRKDVTAGSAYS   89
consensus   *****!*!***********!!!!!!!!!!!!!!!!!!!!!**!******!!!!!*!!!!!!!!!******!!!!!!**!!!!!!!!!*

C15       MT.AGKTNTYSRMRHGYTRNDG...DSTGKKYTRHNGSTYDATKTAVKAKYDVYDDYADDSSHSHYDTDNRVYKSTTARGA.SNRDKHKS  173
G69       MT.AGKTNTYSRMRHGYTRNDG...DSTGKKYTRHNGSTYDATKTAVKAKYDVYDDYADDSSHSHYDTDNRVYKSTTARGA.SNRDKHKS  173
P17       MT.AGKTNTYSRMRHGYTRNDG...DSTGKKYTRHNGSTYDATKTAVKAKYDVYDDYADDSSHSHYDTDNRVYKSTTARGA.SNRDKHKS  173
S10       MT.AGKTNTYSRMRHGYTRNDG...DSTGKKYTRHNGSTYDATKTAVKAKYDVYDDYADDSSHSHYDTDNRVYKSTTARGA.SNRDKHKS  173
S20       MT.AGKTNTYSRMRHGYTRNDG...DSTGKKYTRHNGSTYDATKTAVKAKYDVYDDYADDSSHSHYDTDNRVYKSTTARGA.SNRDKHKS  173
T15       MT.AGKTNTYSRMRHGYTRNDG...DSTGKKYTRHNGSTYDATKTAVKAKYDVYDDYADDSSHSHYDTDNRVYKSTTARGA.SNRDKHKS  173
S40       MT.AGKTNT.YSRMHGYTRDDG...DSTGKKYTRHNGSTYDATKTAVKAKYNVYDDYADDSSHSHYDTDNRVYKSTTARGASNRYTKHKS  173
I12       MT.AGKTNTYSRMRHGYTRDDG...NSTGKKYTRHNGSTYDATKTAVKAKYDVYDDYADDSSHSHYDTDNHVYKSTTARGASNRDTKHKS  173
D15       MT.AGKTNTYSRMRHGYTRDDG...DSTGKKYTRHNGSTYDATKTAVKAKYDVYDDYADDSSHSHYDTDNRVYKSTTARGASNRDTKHKS  174
L42       MT.AGKTNTYSRMRHGYTRDDG...DSTGKKYTRHNGSTYDATKTAVKAKYDVYDDYADDSSHSHYDTDNRVYKSTTARGASNRDTKHKS  174
S26       MTWAGKTNTYSRMRHGYTRDDG...DSTGKKYTRHNGSTYDATKTAVKAKYDVYDDYADDSSHSHYDTDNRVYKSTTARGASNHDTKHKS  175
D43       MT.AGKTNTYSRMRHGYTRDDG...DSTGKKYTRHNGSTYDATKTAVKAKYDVYDDYADDSSHSHYDTDNRVYKSTTARGASNRDTKHKS  174
D49       MT.AGKTNTYSRMRHGYTRDDG...DSTGKKYTRHNGSTYDATKTAVKAKYDVYDDYADDSSHSHYDTDNRVYKSTTARGASNRDTKHKS  174
X15       MT.AGKTNTYSRMRYGYTRDDG...DSTGKKYTRHNGSTYDATKTAVKAKYDVYDDYADDSSHSHYDTDNRVYKSTTARGASNRDTKHKS  174
I27       MT.AGKTNTYSRMRHGYTRDDG...DSTGKKYTRHNGSTYDATKTAVKAKYDVYDDYADDSSHSHYDTDNRVYKSTTARGASNRDTKHKS  174
C01       MT.AGKTNTYSRMRHGYTRDDG...DSTGKKYTRHNGSTYDATKTAVKAKYDVYDDYADDSSHSHYDTDNRVYKSTTARGASNRDTKHKS  174
D11       MT.AGKTNTYSRMRHGYTRDDG...DSTGKKYTRHNGSTYDATKTAVKAKYDVYDDYADDSSHSHYDTDNRVYKSTTARGASNRDTKHKS  174
D48       MT.AGKTNTYSRMRHGYTRDDG...DSTGKKYTRHNGSTYDATKTAVKAKYDVYDDYADDSSHSHYDTDNRVYKSTTARGASNRDTKHKS  174
S11       MT.AGKTNTYSRMRHGYTRDDG...DSTGKKYTRHNGSTYDATKTAVKAKYDVYDDYADDSSHSHYDTDNRVYKSTTARGASNRDTKHKS  174
D13       MD.GDRTHSTYTRMRGYTRSMGDDRDKTGTKYTRHNG.TSYDRKTMRVAKRYVYDDYA.DDGNRHYDKNDVVYKSSTARGAVSDAT.YKG  172
D20       D..GDRTHSTYTRMRGYTRSMG.DDRGTGTKYTRHNG.TSYDRKTMRVAKRYVYDDYA.DDGNRHYDKNDVVYKSSTARGA.VSNATYKG  173
consensus ** ***!********!!!!**!   **!!*!!!!!!!*!****!!***!!**!!!!!!*!****!!!****!!!!*!!!!!*******!*

C15       DYDTNMKASYDNGMARNTHHHYHAWNKKDCKYANYRSKGSVTSKGSSHMGKCYYSGKADDADKVRYN...  240
G69       DYDTNMKASYDNGMARNTHHHYHAWNKKDCKYANYRSKGSVTSKGSSHMGKCYYSGKADDADKVRYN...  240
P17       DYDTNMKASYDNGMARNTHHHYHAWNKKDCKYANYRSKGSVTSKGSSHMGKCYYSGKADDADKVRYN...  240
S10       DYDTNMKASYDNGMARNTHHHYHAWNKKDCKYANYRSKGSVTSKGSSHMGKCYYSGKADDADKVRYN...  240
S20       DYDTNMKASYDNGMARNTHHHYHAWNKKDCKYANYRSKGSVTSKGSSHMGKCYYSGKADDADKVRYN...  240
T15       DYDTNMKASYDNGMARNTHHHYHAWNKKDCKYANYRSKGSVTSKGSSHMGKCYYSGKADDADKVRYN...  240
S40       DYDTNMKASYDNGMARNTHHHYHAWNKKDCKYANYRSKGSVTSKGSSHMGKCYYSGKADDADKVRYN...  240
I12       DYDTNMKASYDNGMARNTHHHYHA.WNKDCKYANYRSKGSVTSKGSSHMGKCYYSGKADDADKVRYN...  239
D15       DYDTNMKASYDNGMARNTHHHYHAWNKKDCKYANYRSKGSVTSKGSNHMGKCYYSGKADDADKTRYN...  241
L42       DYDTNMKASYDNGMARNTHHHYHAWNKKDCKYANYRSKGSVTSKGSNHMGKCYYSGKADDADKVRYN...  241
S26       DYDTNMKASYDNGMARNTHHHYHAWNKKDCKYANYRSKGSVTSKG.SSHMKCYYSGKADDADKVRYSRMN  244
D43       DYDTNMKASYDNGMARNTHHHYHAWNKKDCKYANYRSKGSVTSKG.SSHMKCYYSGKADDADKVRYN...  240
D49       DYDTNMKASYDNGMARNTHHHYHAWNKKDCKYANYRSKGSVTSKG.SSHMKCYYSGKADDADKVRYN...  240
X15       DYDTNMKASYDNGMARNTHHHYHAWNKKDCKYANYRSKGSVTSKG.SSHMKCYYSGKADDADKVRYN...  240
I27       DYDTNMKASYDNGMARNTHHHYHAWNKKDCKYTNYRSKGSVTSKG.SSHMKCYYSGKADDADKVRYN...  240
C01       DYDTNMKASYDNGMARNT.HHYHAWNKKDCKYANYRSKGSVTSKG.SSHMKCYYSGKADDADKVRYN...  239
D11       DYDTNMKASYDNGMARNT.HHYHAWNKKDCKYANYRSKGSVTSKG.SSHMKCYYSGKADDADKVRYN...  239
D48       DYDTNMKASYDNGMARNT.HHYHAWNKKDCKYANYRSKGSVTSKG.SSHMKCYYSGKADDADKVRYN...  239
S11       DYDTNMKASYDNGMARNTHHHYHAWNKKDCKYANYRSKGSVTSKG.SSHMKCYYSGKADDADKVRYN...  240
D13       DYDTNMKASYDNGMARNTRSNVDKKYAYYVVRDDNRTARGRHRNG...........TAYRHHDTKSTYKD  231
D20       DYDTNMKASYDNGMARNTRSNVNKKYAYYVVRDDNRTARGRYRNG...........TAYRHHDTKSTYKD  232
consensus !!!!!!!!!!!!!!!!!!*****************!********!************!*********
```

X| non-conserved
X ≥50% conserved

**Figure S7:** Multiple sequence alignment of *cpsR* (SSU_RS02755) protein sequences. Strains D43 and D49 do not share sequence similarity with other serotype 2 strain, except from a R/S SNP in position 17.

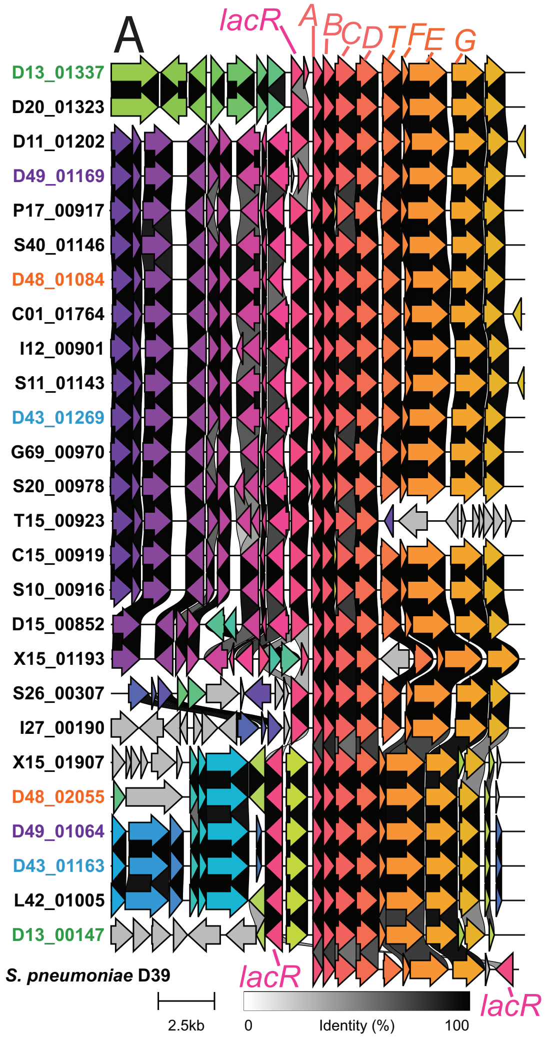


B

| | THY | | | | APS | | | |
|---|---|---|---|---|---|---|---|---|
| | *lacR* | *lacA* | *lacB* | *lacT* | *lacR* | *lacA* | *lacB* | *lacT* |
| C01 | 29.9 | 550.2 | 574.4 | 835.4 | 6.8 | 13.3 | 18.8 | 10.3 |
| C15 | 19.6 | 471.6 | 258.1 | 111.4 | 13.3 | 13.3 | 6.4 | 6.7 |
| D11 | 118.5 | 69.2 | 126.9 | 122.9 | 7.5 | 0.8 | 0.2 | 5.1 |
| D13 | 418.8 | 265.0 | 321.3 | 51.7 | 508.4 | 208.1 | 280.2 | 95.3 |
| D15 | 13.8 | 226.9 | 571.0 | 184.6 | 7.5 | 3.3 | 3.3 | 6.6 |
| D20 | 137.1 | 1264.1 | 1352.5 | 346.5 | 59.1 | 2.0 | 2.2 | 5.5 |
| D43 | 122.3 | 218.9 | 89.4 | 60.5 | 25.8 | 23.6 | 17.3 | 4.0 |
| D48 | 35.7 | 21.1 | 20.5 | 11.3 | 46.5 | 25.9 | 16.3 | 5.3 |
| D49 | 74.4 | 34.3 | 32.8 | 3.1 | 107.5 | 11.3 | 9.1 | 3.8 |
| G69 | 43.3 | 502.3 | 319.7 | 230.1 | 10.9 | 19.6 | 12.7 | 8.1 |
| I12 | 13.6 | 9.8 | 19.2 | 9.3 | 16.5 | 9.5 | 27.2 | 11.0 |
| I27 | 195.8 | 1021.7 | 2307.4 | 1555.6 | 208.0 | 6.4 | 7.4 | 6.8 |
| L42 | 98.4 | 273.1 | 196.6 | NA | 14.3 | 7.4 | 6.1 | NA |
| P17 | 9.0 | 19.1 | 17.3 | 49.4 | 10.8 | 13.6 | 10.5 | 4.2 |
| S10 | 38.8 | 452.8 | 314.8 | 191.9 | 8.5 | 13.1 | 11.7 | 5.2 |
| S11 | | | | | 6.8 | 28.6 | 56.5 | 16.8 |
| S20 | 60.9 | 738.0 | 562.5 | 247.7 | 7.0 | 28.0 | 16.4 | 5.2 |
| S26 | 195.2 | 201.9 | 539.0 | 300.7 | 83.2 | 6.0 | 9.9 | 17.8 |
| S40 | 9.3 | 3.7 | 8.4 | 10.2 | 11.6 | 9.0 | 9.1 | 6.1 |
| T15 | 21.5 | 107.6 | 105.6 | 31.2 | 8.1 | 6.4 | 6.6 | 5.7 |
| X15 | 74.2 | 140.9 | 134.1 | 3.4 | 44.3 | 19.1 | 19.2 | 1.8 |

**Figure S8:** Divergent *lac* gene cluster expression in strain D13. **A)** Several variants of the *lac* gene cluster are carried by the strains, and these may occur alone or two (at different locations) in the same genome. D13 has 2 *lac* gene clusters, which both feature inverted or pseudogene *lacR* copies (although other strains sharing the inverted variant had exopressino as expected). Genomes with several copies highlighted in color. Annotation based on Afzal et. al. 2014 (https://doi.org/10.1128/AEM.01370-14) and *S. pneumoniae* D39 assembly CP000410.2. **B)** *lacR* is highly expressed in D13, but *lacA* and *lacB* (and *lacT)* are nonetheless highly expressed in both media. P1/7 has unexpectly low *lac* expression compared to other ST1 strains.

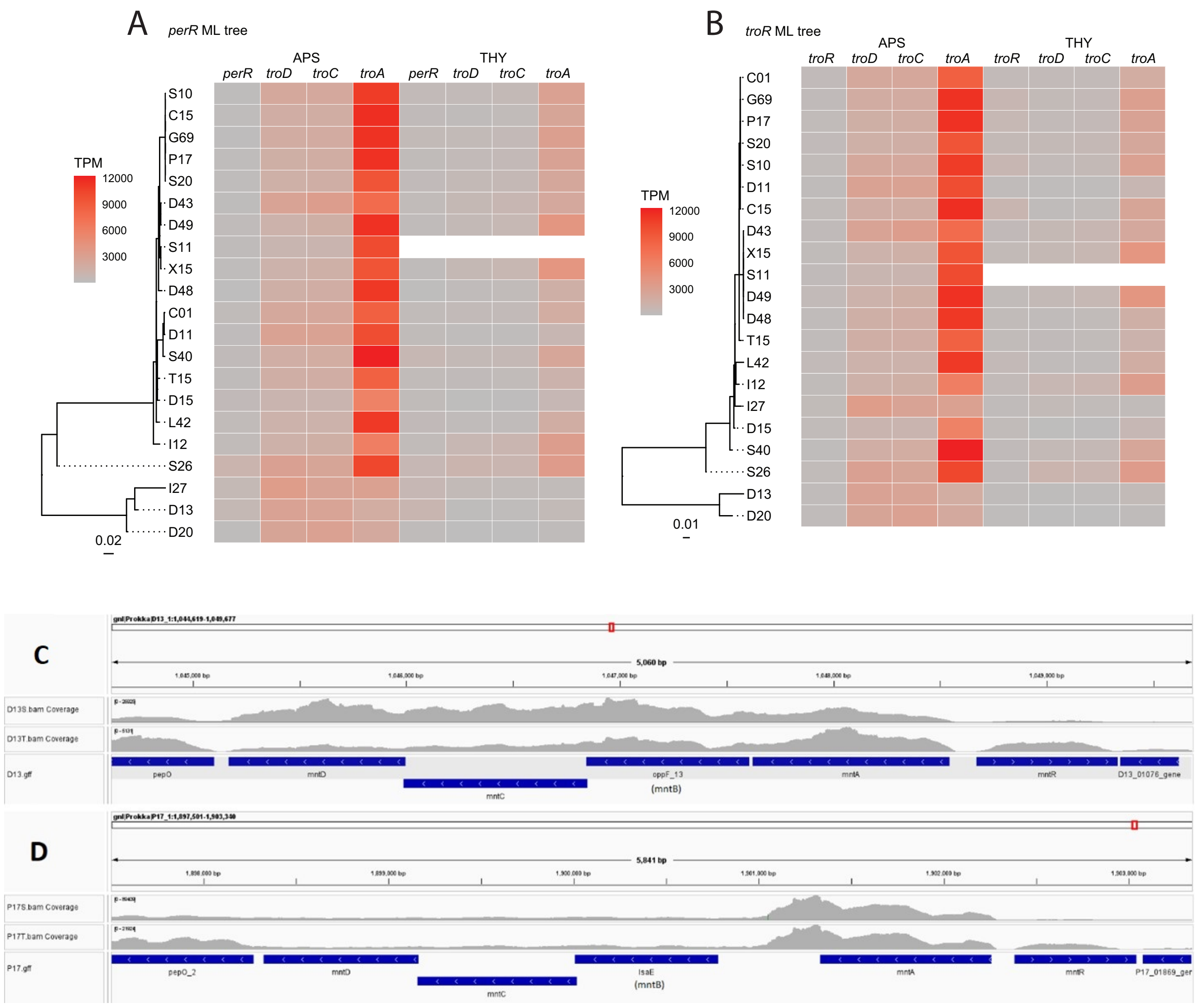


**Figure S9:** Gene expression of the *tro*/*mnt* gene cluster. A) ML tree of *perR* with *tro* expression heatmap. B) ML tree of *troR* with *tro* expression heatmap. C) Raw RNA-seq coverage of the *tro* gene cluster in strain D13 (visualized in IGV, https://igv.org). D) Raw RNA-seq coverage of the *tro* gene cluster in strain P1/7.
